# Cost-effectiveness of quadrivalent versus trivalent inactivated influenza vaccines for the Portuguese elderly population

**DOI:** 10.1101/2022.01.04.474923

**Authors:** Diana Tavares, Helena Mouriño, Cristina Antón Rodríguez, Carlos Martín Saborido

## Abstract

**Background:** Quadrivalent Inactivated Vaccine (QIV) is expected to replace Trivalent Inactivated Vaccine (TIV) over time. In Portugal, TIV is free of charge for risk groups, which include older adults. In its turn, QIV – which provides broader protection as it includes an additional lineage B strain – was introduced in Portugal in October 2018, but only from the influenza season 2019/20 was provided free of charge for the risk groups. This study evaluates the cost-effectiveness of switching from TIV to QIV from the National Health Service perspective in the Portuguese elderly mainland population (≥ 65 years old).

**Methods:** A decision tree model was developed to compare TIV and QIV, based on Portuguese hospitalization data for the 2015/16 influenza season. The primary health economic outcome under consideration was the Incremental Cost-Effectiveness Ratio (ICER). In addition, a one-Way Sensitivity Analysis was performed to evaluate the impact of model parameters on the ICER; Probabilistic Sensitivity Analysis was also carried out to analyze the robustness of the base case results.

**Results:** The high cost of QIV (approximately three times the cost of TIV) would lead to a total increment of 5,283,047 €, and the resulting ICER would be 26,403,007€/QALY, mainly above the usual willingness-to-pay threshold.

**Conclusions:** From the National Health Service perspective, our findings reveal that QIV is not cost-effective for the Portuguese elderly population due to the high cost. If the QIV costs were the same as the TIV, then QIV would be cost-effective.

## Introduction

Seasonal influenza is an acute respiratory disease caused by infection with influenza viruses(1). There are four types of seasonal influenza viruses: A, B, C and D(1). Only influenza A and B viruses circulate and cause seasonal epidemics of disease(1). The infection may cause signs and symptoms like fever, cough, headache, muscle and joint pain and weakness(1).

Influenza outbreaks are recorded every year. In temperate zones of the northern and southern hemispheres, epidemics occur during the winter, while in the tropics they occur throughout the year(2,3). In Portugal, influenza outbreaks are characterized by significant morbidity in the general population and increased mortality rates. The high-risk groups are: elderly (≥65 years old), patients with chronic or immunosuppressive conditions aged six months or older, pregnant women, health professionals and other healthcare givers(4).

An influenza pandemic can appear when a new and different type of influenza A virus emerges, and it can infect humans who are not immunized(1,5). The most recent pandemic was in 2009, caused by the A(H1N1)2009 virus, after which it became a seasonal influenza serotype(1). In Portugal, 1,189 people were hospitalized, around 10% in intensive care units, resulting in 124 deaths(6).

According to the World Health Organization (WHO), vaccination is the most effective way to prevent seasonal influenza and subsequent severe outcomes(1). Therefore, vaccination should be administered annually to provide optimal protection. Influenza Surveillance System in Portugal is usually activated on week 40 (October) of a given year (*n*) and lasts up to week 20 (April) of the following year (*n*+1) (7,8). Thus, it is recommended to administer the vaccine during autumn and winter, preferably until the end of the year *n* (4). The development of immunity against influenza viruses takes about two weeks after vaccination(9). However, vaccine effectiveness depends on the influenza subtypes included in the vaccines (a mismatch occurs when the viruses in the vaccine are different from the circulating viruses) and the population vaccination coverage. Unfortunately, several European countries reported a decline in vaccination coverage among older people from 2008/09 to 2014/15 seasons. Conversely, Portugal reported an increase in vaccination uptake throughout these seasons(10). In the 2015/16 season, influenza vaccination coverage was 50.1% in Portugal(7). The coverage rate has still increased in the last few years. For instance, in the 2017/18 season, it has increased to 60.8% for people aged 65 years or above(11).

A Cochrane study on this topic concluded that the risk of influenza decreases from 6% to 2.5%, and the risk of Influenza-Like Illness (ILI) reduces from 6% to 3.5%, between unvaccinated and vaccinated groups (≥ 65 years old) during a single season(12). In the European Union, it is estimated that influenza vaccination prevents up to 37,000 deaths each year(13).

There are two main types of seasonal influenza vaccines, namely Inactivated Influenza Vaccine (IIV) and Lived Attenuated Influenza Vaccine (this one is not within the scope of the paper)(5,12). The different IIV developed are: trivalent, trivalent adjuvanted and quadrivalent. Trivalent Inactivated Vaccine (TIV) is the most common IIV and protect against three different influenza viruses (two influenza A strains and one influenza B lineage). In some EU/EEA countries, the trivalent adjuvanted vaccine is available for older people to empower immune response(5,14). Quadrivalent inactivated Influenza Vaccine (QIV) is available since the 2014/2015 season in some EU/EEA countries, and it is expected to replace TIV over time(5). It contains one more influenza B strain in addition to those included in TIV, which provides wider protection against influenza viruses. TIV is available in all EU/EEA countries, while QIV is only available in some of them. In Portugal, QIV was launched in October 2018, but only from the influenza season 2019/20 it was provided free of charge for the risk groups.

Influenza is, thus, a major global health threat that affects all countries, not only from a health perspective but also from economic and social approaches. Therefore, it is crucial to carry out economic evaluations of the influenza vaccination programmes.

This work aims to estimate the cost-effectiveness of switching from TIV to QIV based on the National Health Service perspective. The results are presented as incremental cost per unit of effect, described by the Incremental Cost-Effectiveness Ratio (ICER). As far as we know, no cost-effectiveness analysis of Portuguese influenza vaccination has been reported in the literature yet.

## Methods

### Key Features of the Economic Evaluation

The target population is the population of mainland Portugal aged 65 and above because they are one of the most relevant influenza risk groups. In 2015, they represented 21% of the Portuguese population(15). This group is usually retired, so productivity losses are not relevant. The cost-effectiveness evaluation was performed from the National Health Service (NHS) perspective, where only direct costs were considered. Thus, only the medical costs paid by the NHS are considered here.

The Influenza Surveillance System runs from week 40 of a specific year until week 20 of the following year, so the time horizon established for this work was one year. Therefore, data under analysis refers to the 2015/16 influenza season.

As stated above, economic evaluations are performed to identify, measure, value and compare the costs and consequences of different therapeutic alternatives in what respects to influenza vaccination. To perform the cost-effectiveness analysis, health outcomes generated by the two alternative interventions under analysis are assessed through the Quality-Adjusted Life Year (QALY). It measures the length of life-years (i.e., quantity gains) and the quality of life (i.e., quality gains). The quality of life is quantified by the notion of utility: the greater the preference for a particular state, the greater its utility. Utilities incorporate in the analysis the preferences of individuals for different treatment-related outcomes. Another key measure is the concept of quality-of-life loss due to the disease (influenza, in this case), that is, disutility. It corresponds to the reverse of utility, i.e., decrease in utility due to serious health complications. These quantities are obtained by Health-related quality of life (HRQoL) instruments, such as the EQ-5D family of questionnaires. Studies about utilities and disutilities for the Portuguese population are very scarce. In this work, utilities associated with different health states were obtained from a study on *EQ-5D Portuguese Population norms*(16). Data available were stratified by age, but unfortunately, there was no information for people aged 65 or over. Therefore, the utilities for that age group were derived by interpolation (see S1 Appendix). Moreover, no Portuguese studies on disutilities were found for influenza. Such data were extracted from a Spanish study (17) (see S1 Appendix). All these results were thus combined in a single index, QALY(18,19).

Incremental Cost-Effectiveness Ratio (ICER) is the primary cost-effectiveness outcome used in this work(19). To make recommendations to policymakers, ICER is usually compared with an established threshold, called Willingness-To-Pay (WTP). Some decision-makers have established a Willingness-To-Pay (WTP) threshold, that refers to the maximum ICER accepted for an intervention to be considered cost-effective. In Portugal, there is no established cost-effectiveness threshold for health interventions. Some Portuguese authors refer to a WTP threshold of 30,000€/QALY(20,21), while others consider the cost-effectiveness threshold twice the Gross Domestic Product (GDP) per capita or Gross National Income (GNI) per capita(21,22). In 2015, the Portuguese GNI per capita was 16,887.36€, resulting in a ceiling ratio of approximately 34,000€ (twice the GNI)(23,24). In this paper, the ICER will be used to provide information on the extra amount that is necessary to pay in order to gain an extra QALY when the most effective alternative is chosen(19,25).

### Decision Tree

The cost-effectiveness analysis was based on a decision tree model, a very powerful tool in this context because it incorporates probabilities of the outcomes, resource costs, vaccination coverage rate, prices of the vaccines, efficacy, utilities and disutilities. It consists of a series of pathways representing possible prognoses for each alternative therapy being evaluated. A static model was used, and no herd effects of vaccination were evaluated.

One decision tree model was developed for each alternative therapy (TIV and QIV). For convenience, each decision tree was split into two different diagrams referring to vaccinated and unvaccinated people. For illustration, Fig 1 displays the branch of the decision tree for vaccinated individuals.

ILI, influenza-like-illness; H, healthy; D, death; Conf. Influenza, Confirmed Influenza; GP Consultation, General Practitioner Consultation; Pneu. Hosp., Hospitalization due to pneumonia; RD Hosp., Hospitalization due to respiratory disease; HD Hosp., Hospitalization due to heart disease; Inf. Hosp., Hospitalization due to Influenza.

**Fig 1.**
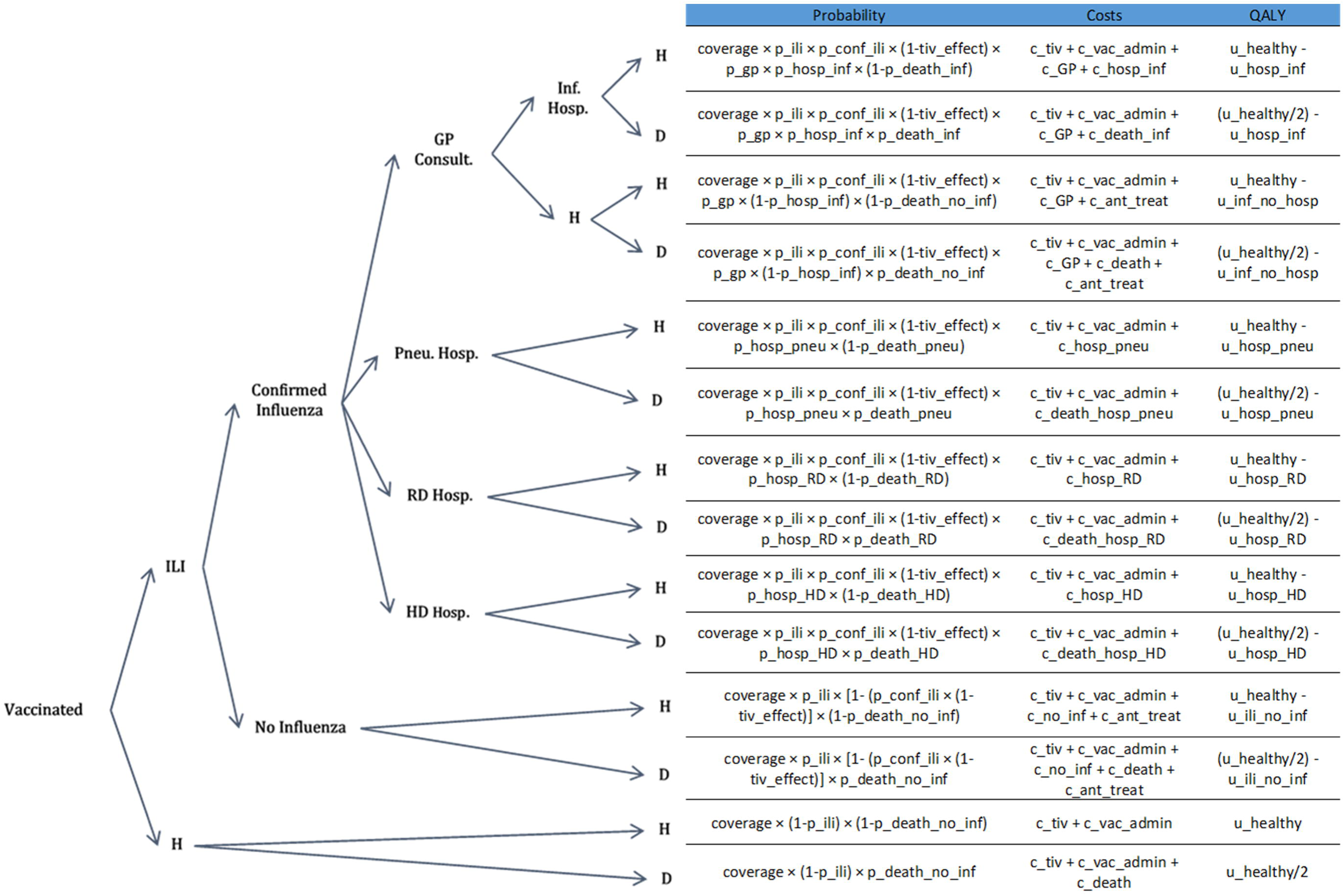
Decision tree for the cost-effectiveness analysis. Branch for the vaccinated people aged ≥ 65 years old.

It was assumed that influenza confirmation (through diagnostic tests) was carried out in the context of General Practitioner (GP) consultation, pneumonia hospitalization, respiratory disease hospitalization or heart disease hospitalization. Although it is known that influenza may be diagnosed in many other situations, the diseases described above are usually associated with complications due to influenza in the elderly. It should be noted that influenza event (as a secondary diagnosis) may have occurred before or after hospital admission, in which principal diagnoses was pneumonia, respiratory disease or heart disease. It is important to note that the “Healthy” state is assumed to include all individuals who did not die, regardless of their health conditions.

### Input Parameters and Computation of the Probabilities

The decision tree is powered by different types of parameters. A detailed description of each of these measures is provided in the S2 Appendix (which includes S1 Table). A summary of the base case values for the parameters under consideration is displayed in Table 1. The computation of the different parameters involved in the decision tree pathways for the TIV – probabilities, costs and QALY – are given in the S1 Fig and S2 Table. Similar calculations are made for the QIV scenario.

**Table 1.**
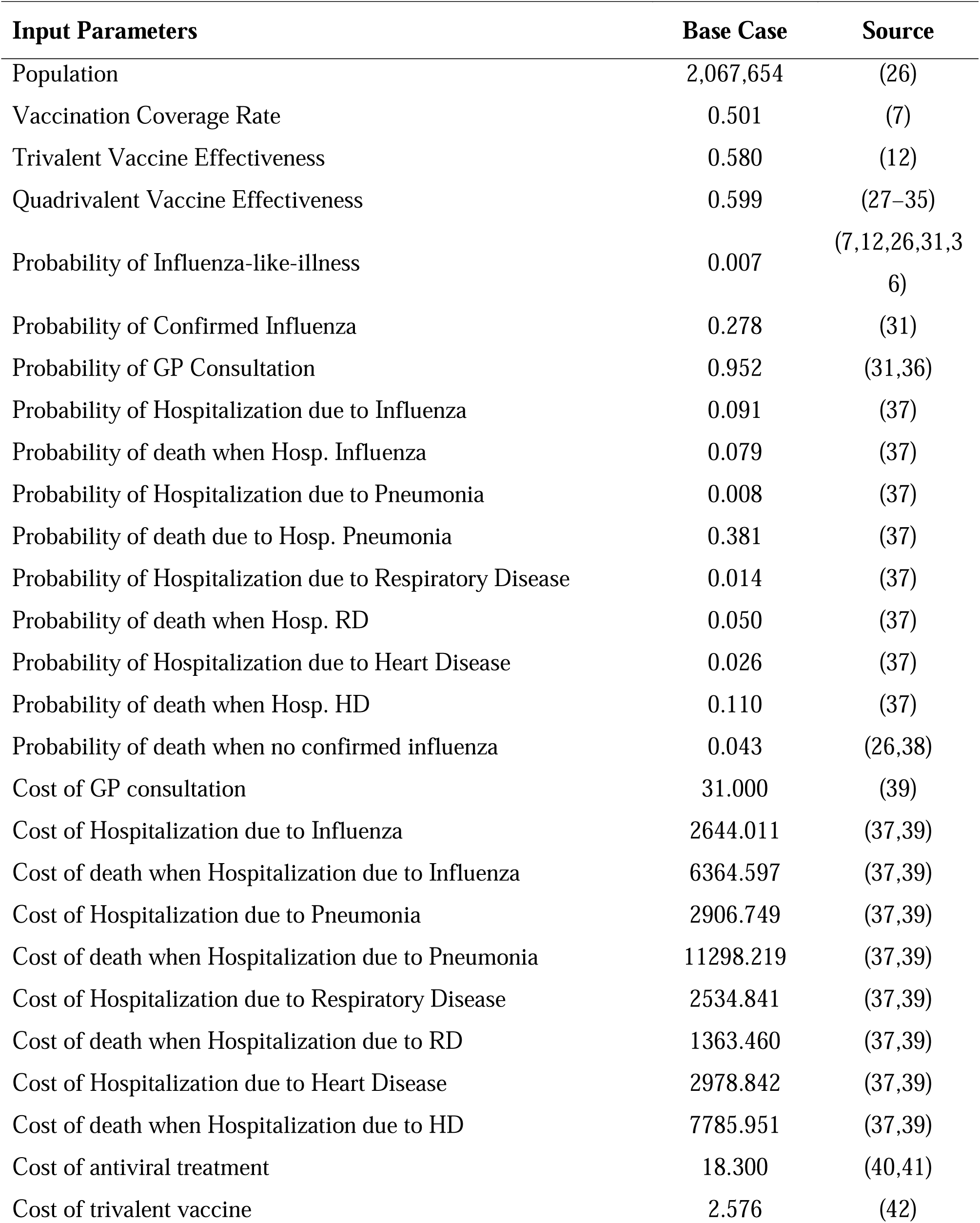
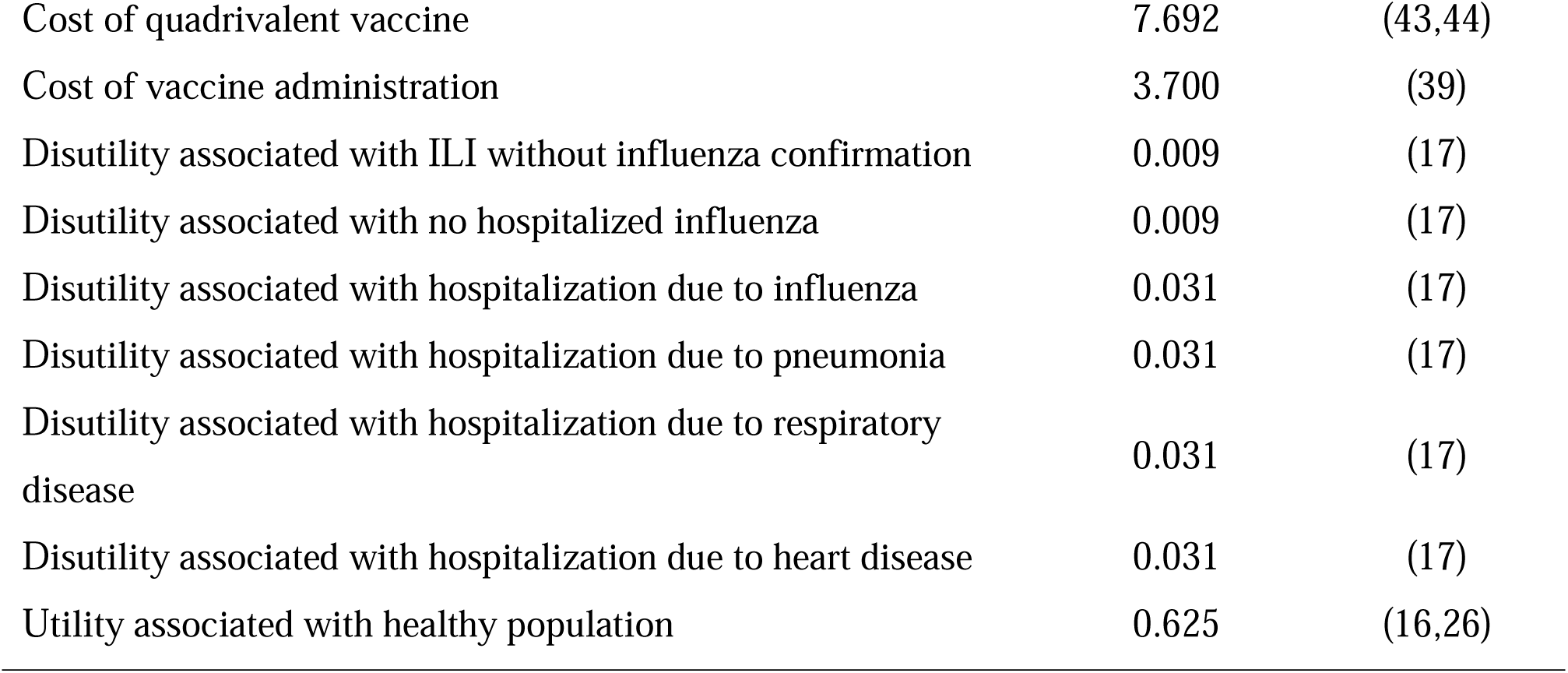
Input parameters for the decision tree. Base case values.

### Scenario Analysis

The most common ways to characterize parameter uncertainty are: One-way Sensitivity Analysis (OWSA) and Probabilistic Sensitivity Analysis (PSA)(19,25).

OWSA consists of varying the point estimates of the input parameters one at a time within a plausible range of values and assessing the impact on model outcomes(19,25), which corresponds to the ICER. When the Confidence Interval (CI) for a specific parameter was known from the literature, these limits were used as the OWSA range. For parameters with no CI available, a range of ±20% was used to assess model’s sensitivity to parameters variation. A tornado diagram was used to report OWSA results.

A scenario analysis of the vaccination coverage rate, TIV effectiveness and QIV effectiveness variation was also carried out. The cost of the quadrivalent vaccine was also studied in detail, as it was identified as a critical parameter(45). For vaccination coverage rate and QIV cost, a sequence of parameters values was generated with an increment of 0.05. For TIV and QIV effectiveness, the increment used was 0.005.

Instead of representing the parameters as single point estimates as in OWSA, in PSA parameters are represented by random variables following a particular distribution. The S2 Table displays the base case values used for each input parameter of the decision tree models. It also shows the probability distribution (with information on location, shape, scale, when appropriate) used for each parameter under consideration. A total of 1,000 Monte Carlo (MC) simulations were performed.

## Results

### Base Case Analysis

Table 2 summarises the base case results of shifting from TIV to QIV in the elderly, based on the 2015/16 influenza season data. Results show a difference in confirmed influenza cases of approximately 37. As a result, about 36 GP consultations could have been averted and, therefore, 1,103€ saved. In addition, five hospitalizations, including hospitalizations due to influenza, pneumonia, respiratory disease and heart disease, could have been averted and one life saved.

**Table 2.** Base Case. Results of the cost-effectiveness evaluation comparing TIV and QIV.

|  | TIV | QIV | Difference<br>(QIV-TIV) |
| --- | --- | --- | --- |
| Events |  |  |  |
| GP Consultations | 2631.35 | 2595.77 | -35.58 |
| Hospitalizations due to Influenza | 239.00 | 235.77 | -3.23 |
| Deaths due to Influenza Hospitalization | 19.00 | 18.74 | -0.26 |
| Hospitalizations due to Pneumonia | 21.00 | 20.72 | -0.28 |
| Deaths due to Pneumonia Hospitalization | 8.00 | 7.89 | -0.11 |
| Hospitalizations due to RD | 40.00 | 39.46 | -0.54 |
| Deaths due to RD Hospitalization | 2.00 | 1.97 | -0.03 |
| Hospitalizations due to HD | 73.00 | 72.01 | -0.99 |
| Deaths due to HD Hospitalization | 8.00 | 7.89 | -0.11 |
| Vaccine Doses | 1,035,895 | 1,035,895 | 0.00 |
| Costs |  |  |  |
| GP Consultations | 81,572 € | 80,469 € | -1,103 € |
| Hospitalizations due to Influenza | 710,464 € | 701,189 € | -9,275 € |
| Deaths due to Influenza Hospitalization | 121,552 € | 119,934 € | -1,617 € |
| Hospitalizations due to Pneumonia | 128,213 € | 126,508 € | -1,705 € |
| Deaths due to Pneumonia Hospitalization | 90,401 € | 89,189 € | -1,211 € |
| Hospitalizations due to RD | 99,125 € | 97,841 € | -1,285 € |
| Deaths due to RD Hospitalization | 2,731 € | 2,697 € | -34 € |
| Hospitalizations due to HD | 256,048 € | 252,687 € | -3,361 € |
| Deaths due to HD Hospitalization | 62,303 € | 61,471 € | -831 € |
| Vaccine Doses | 2,668,465 € | 7,968,557 € | 5,300,092 € |
| Total | 8,018,570 € | 13,301,618 € | 5,283,047 € |
| QALYs |  |  |  |
| Total | 1265456.95 | 1265457.15 | 0.20 |
| ICER (€/QALY) |  |  | 26,403,007 |

Regarding hospitalization and death costs, caution must be taken in the interpretation of the results. The cost of death, when hospitalized, includes all hospitalization costs, and in the same way, the hospitalization costs also include the costs of people who died. Thus, 15,625€ could have been saved in hospitalizations: 9,275 € in hospitalizations due to influenza; 1,705 € in hospitalizations due to pneumonia; 1,285€ in hospitalizations due to respiratory disease; and 3,361 € in hospitalizations due to heart disease. From that value, 3,694€ are related to hospitalizations of patients who died (Table 2).

In what concerns the number of vaccinated people (mentioned in Table 2 as vaccine doses), the value is the same for both strategies. It was an expected result because the vaccination coverage rate applied to TIV and QIV models was the same (the paper’s focus is to evaluate the cost-effectiveness of shifting from TIV to QIV). However, there is a difference in the expected costs computed from the two strategies because they depend on the cost of each vaccine. A total of 1,035,895 vaccine doses were expected to have been administered to individuals aged ≥65 years old in the 2015/16 season. For the TIV strategy, this represented spending 2,668,465 €, while for QIV corresponds to 7,968,557 €, that is, shifting from TIV to QIV would result in an additional cost of 5,300,092 € (Table 2).

The expected difference in costs between TIV and QIV is 5,283,047 €, while the difference in effects (QALYs) is expected to be 0.20. These values result in a base case ICER of 26,403,007€/QALY (Table 2). The plot of the differences in costs and effects on the cost-effectiveness plane shows the value under consideration is in the first quadrant. Thus, for a cost-effectiveness threshold of 34 000€/QALY (twice the GNI per capita), the new vaccine would not be cost-effective.

### One-way Sensitivity Analysis

Fig 2 summarizes the one-way sensitivity analysis using the tornado diagram.

**Fig 2.**
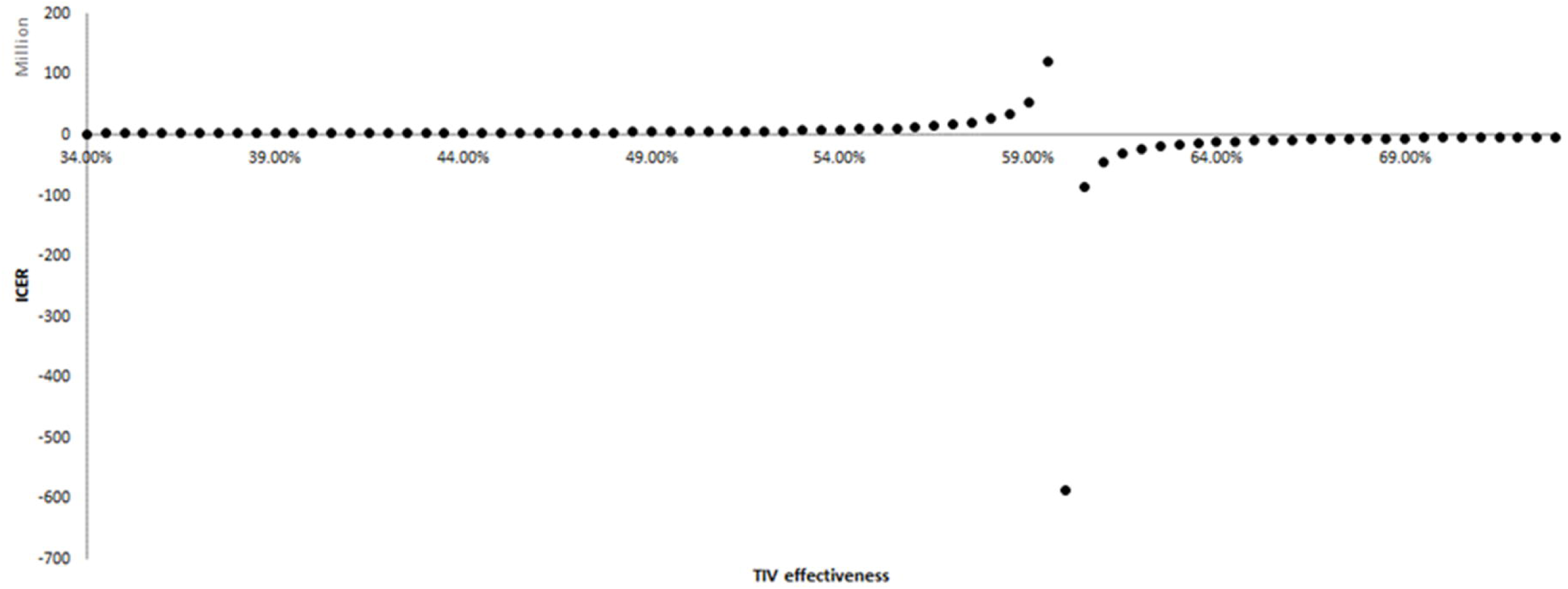
One-way sensitivity analysis. Tornado diagram.

Fig 2 reveals that the disutility associated with ILI when no confirmed influenza, and the disutility associated with no hospitalized influenza have the highest impact on the ICER. For example, if the first parameter mentioned above assumes the value of 0.007 (lower bound), it results in an ICER of 19,219,556 €/QALY; and the value of 0.011 (upper bound) results in an ICER of 42,160,989 €/QALY. This can be explained by the increase in incremental QALYs, yielded by decreasing this parameter, resulting in a reduced ICER and vice-versa. In contrast, the variation of the second parameter within the 95% CI (0.007, 0.011) produced ICERs of 39,019,821€/QALY and 19,951,738€/QALY, respectively. Thus, while the pathway corresponding to vaccinated patients with ILI but not confirmed influenza is more populated in the QIV model than in TIV, the same is not valid for the pathway referring to vaccinated patients with no hospitalized influenza. Thus, a reduction in disutility associated with no hospitalized influenza would decrease incremental QALYs and increase ICER.

The cost of the quadrivalent vaccine is the third parameter appearing in the tornado diagram (Fig 2). The variation of the cost of the quadrivalent vaccine within a ±20% range results in an ICER of 18,438,140 €/QALY (lower bound) and 34,367,874€/QALY (upper bound), respectively. The high impact of QIV’s cost on ICER was expected if attention is paid to the difference in costs of QIV and TIV vaccine doses (Table 2). The cost of QIV’s vaccines doses is largely above the cost of TIV’s vaccine doses. Further analysis was performed, varying the QIV cost within a broader range (Fig 3). For a QIV cost equal to TIV cost, an ICER of -85,182€/QALY was obtained, meaning that shifting from TIV to QIV would be cost-saving, as expected. However, a value higher than 2.576€ would quickly lead to being not cost-effective.

**Fig 3.**
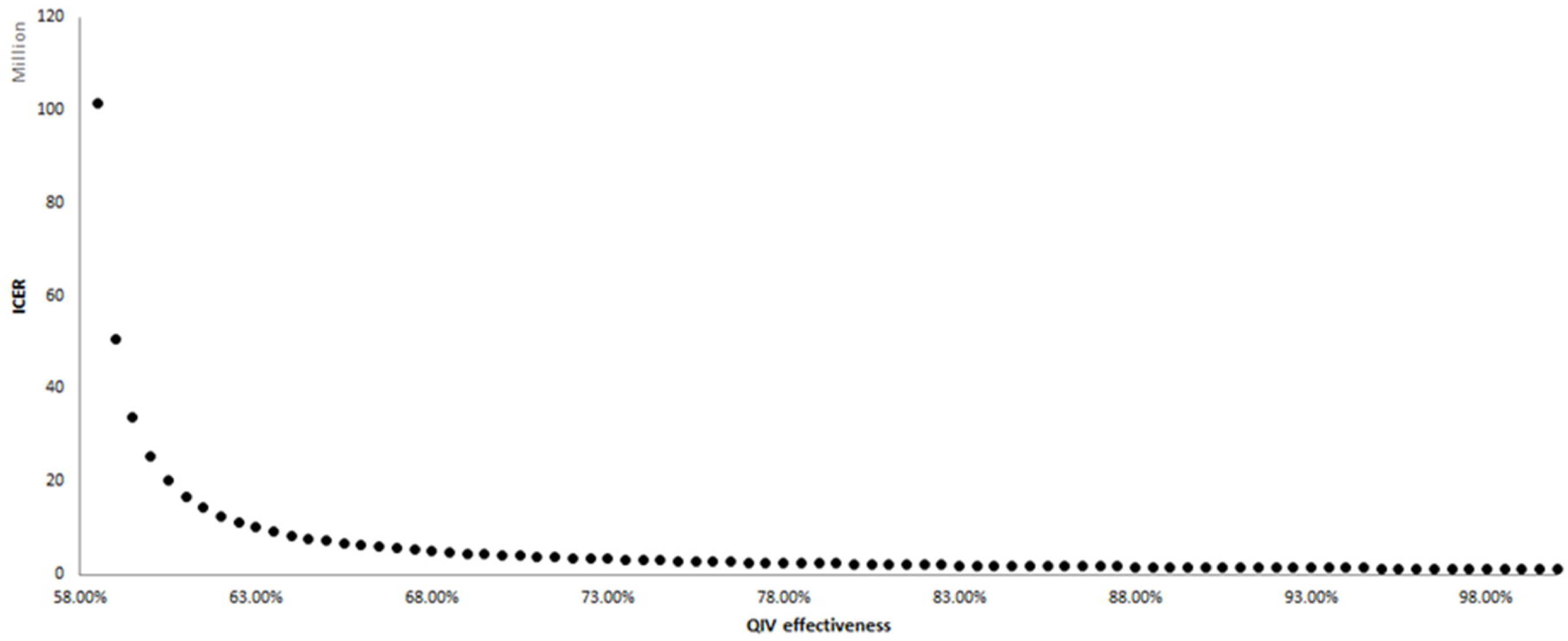
One-way sensitivity analysis of the cost of QIV. Variation between 2.576€ (the cost of TIV) and 9.756€ with increments of 0.05€.

The probability of ILI and the probability of confirmed influenza also have a high impact on ICER (Fig 2). Varying these parameters 20% downward and upward produced ICERs of 33,025,054€/QALY and 21,988,309€/QALY, respectively. Thus, as the probability of ILI and/or the probability of confirmed influenza increases, the ICER decreases.

One-way sensitivity analyses of the effectiveness of QIV and TIV are displayed in S2 Fig and S3 Fig, respectively. If the TIV effectiveness is equal to QIV effectiveness, the ICER is not defined as it results in a difference in effects equal to zero and, therefore, a division by zero, and a vertical asymptote at 59.91% is shown (S2 Fig). As TIV effectiveness approaches 59.91% from the left, the ICER tends to positive infinity. Otherwise, as TIV effectiveness approaches 59.91% from the right, the ICER tends to negative infinity. Such behavior is explained by the difference in effects close to zero when TIV and QIV effectiveness are close. A negative ICER is produced for TIV effectiveness values higher than QIV effectiveness as the incremental QALYs value is negative. Similarly, S3 Fig shows a vertical asymptote when QIV effectiveness equals 58%; ICER tends to infinity when it approaches the vertical asymptote from the right. Finally, the higher the effectiveness of quadrivalent vaccines (>58%), the lower the ICER.

Although vaccination coverage rate does not significantly impact ICER as the above parameters do, a detailed study was also performed for this parameter (S4 Fig). A higher coverage rate would result in decreased ICER. As recommended by the World Health Assembly, for a coverage rate of 75%, 21,010,680 €/QALY would be obtained, corresponding to a reduction of 20,42%.

### Probabilistic Sensitivity Analysis

Probabilistic sensitivity analysis allows one to evaluate the robustness of the base case results. To attain this goal, a probability distribution was assigned to each parameter of the model to reflect parameter uncertainty due to sampling errors. Parameters values were then sampled 1,000 times; expected costs and QALY were recorded from the 1,000 MC simulations. Finally, some empirical statistics such as mean, standard deviation and confidence intervals were computed.

In general, the results from PSA (S3 Table) are similar to those obtained from the base case analysis (Table 2). The estimated mean costs difference between QIV and TIV is 5,293,456 € with an interval estimate of (4,375,217€; 6,088,637€), while the estimated mean effect difference corresponds to 0.20 QALYs with an interval varying from 0.06 to 0.43 QALYs. On its turn, mean ICER is estimated at 34,501,793€/QALY (13,052,665€/QALY; 87,418,632€/QALY). This value is about 31% higher than in the base case result.

S5 Fig displays the scatterplot of the 1,000 MC simulations to compare TIV and QIV, that is, the graphical representation of costs against QALYs. The base case results are also shown. QALY of both interventions varies within a similar range of values (it varies between 1,155,190 and 1,375,507). On the other hand, the difference between TIV and QIV in terms of costs is very well delimited, being QIV more costly than TIV (QIV: mean=13,323,148€; SD=987,347€; TIV: mean=8,029,692€; SD=548,739€).

Fig 4 shows the results of the simulations in a cost-effectiveness plane, where the incremental costs are plotted against the incremental effects. According to Fig 4, the cost-effectiveness results are robustly located in the first quadrant of the cost-effectiveness plane, and all simulated ICERs are higher than CE thresholds previously established.

**Fig 4.**
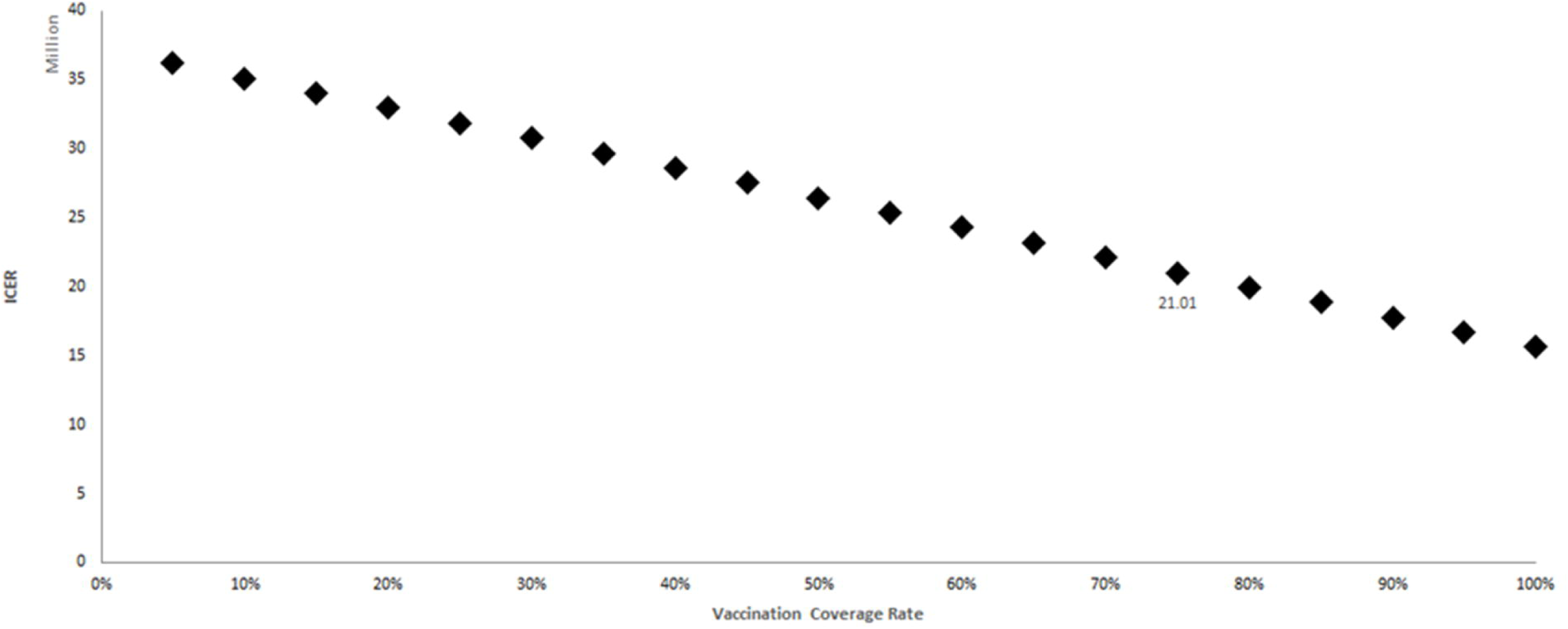
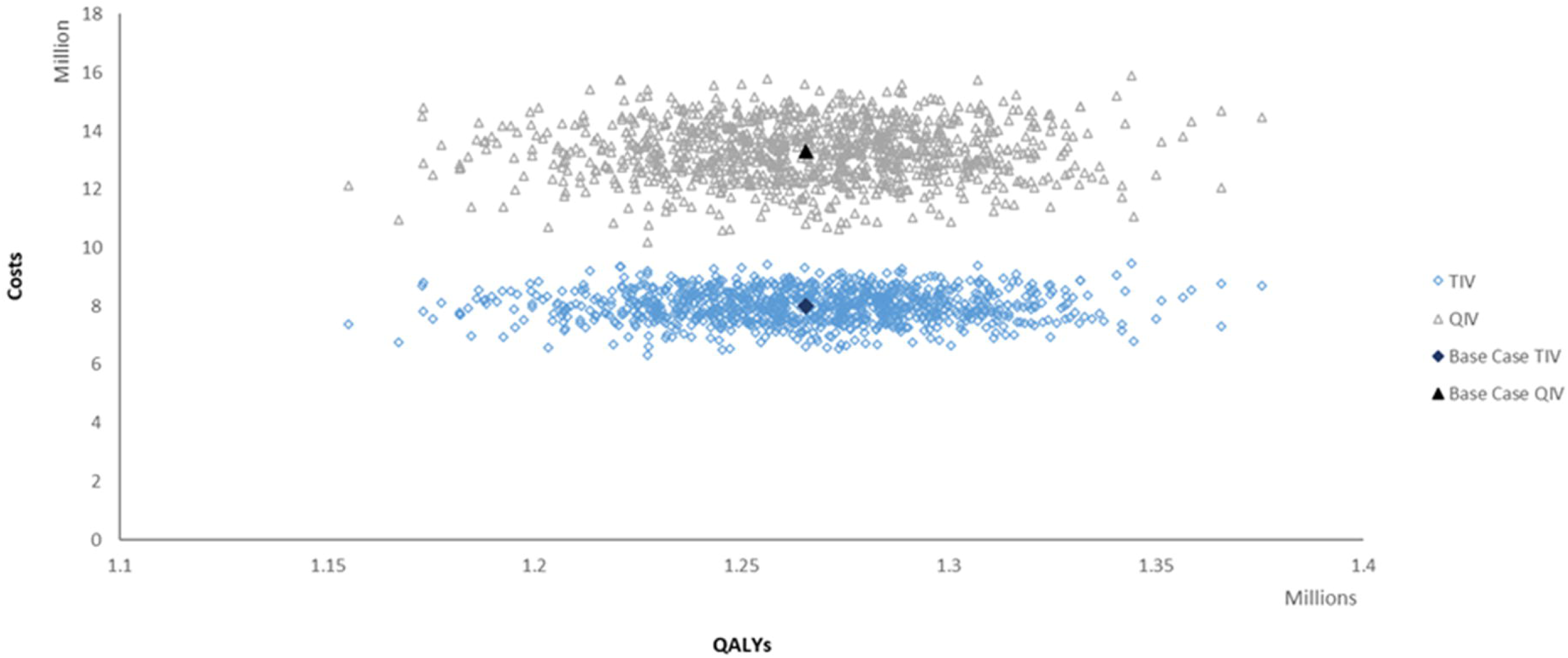
Probabilistic sensitivity analysis for TIV and QIV. Cost-effectiveness plane based on data from the 2015/16 influenza season.

## Discussion

To the best of our knowledge, this is one of the first published studies performing a cost-effectiveness evaluation of seasonal influenza vaccination in Portugal. Only a few papers studied the burden of disease and vaccination effectiveness in the Portuguese population, for instance, the international I-MOVE+ project(46–50).

We evaluated the cost-effectiveness of switching from TIV to QIV based on the 2015/16 influenza season data. Base case results revealed that the universal substitution of TIV with QIV would result in an ICER of 26,403,007€ per QALY gained. As such, for the cost-effectiveness thresholds of 30,000€/QALY or 34,000€/QALY, QIV is not cost-effective. Probabilistic sensitivity analysis enhanced the robustness of the base case results. The ICER is much higher than any possible ceiling ratio established by NHS. Such results are comparable to those obtained for the first year of QIV administration to Hong Kong elderly (≥80 years old) when a cost-effectiveness analysis over nine seasons was performed(51). The need for a longer time horizon study is here emphasized.

The majority of the published international papers(45,52–60)does not report such a high ICER, and QIV is usually identified as cost-effective compared with TIV. As cited above, one of the reasons might be using of a long time horizon to calculate QIV effectiveness, which allowed to assume a few seasons that match the TIV with the circulating strains and therefore low mismatching (see S2 Appendix). Another reason might be the cost of QIV, which in this study is about three times the TIV. Regarding the quadrivalent vaccine, its cost is here 200% higher than TIV. However, in the literature, the additional cost of QIV is rarely higher than 100% of the TIV cost(52,54–56), which might be one of the reasons why QIV is cost-effective in some studies. To be cost-effective, the cost of QIV would need to be the same as TIV in our case.

In the literature, the probability of influenza-like-illness and the probability of confirmed influenza was found to be higher than those used here. For example, Capri et al.(60) applied a probability of influenza of 6.4%, resulting from the product of ILI attack rate (16.8%) by influenza virus isolation rate (32.1%), while Van Bellinghen et al.(55) used a probability of symptomatic influenza infection of 6.17%. In our study, the probability of confirmed influenza was derived from a Portuguese report on influenza surveillance, and the probability of ILI was determined from other input values. It is comparable to the value reported by INSA for the 2015/16 influenza season(31).

One-way sensitivity analysis demonstrated that our model is more sensitive to disutility associated with ILI, disutility associated with no hospitalized influenza, cost of quadrivalent vaccine and probabilities of confirmed influenza and ILI. The most relevant input parameters obtained from the OWSA are in line with other OWSA results in the literature(52,55,56). Considering the PSA, the results are in line with the base case conclusions and confirm their robustness.

A limitation of our study has to do with the fact that the model is static in a way that herd effects of immunization are not considered, and only direct protection is captured. Thus, the impact of vaccination on the burden of the disease might be underestimated. In the future, a dynamic transmission model might be a possible approach to account for the impact of vaccination on influenza transmission(45,56). In addition, side effects of vaccination were not included in this study, but QIV safety is assumed to be comparable to that of TIV, and side effects are assumed to be mild and temporary. Thus, no impact is expected on the ICER(56).

Summing up, this study contributes to understanding the impact of annual influenza epidemics on health economics and public health in Portugal.

## Acknowledgments

H. Mouriño was supported by the National Funding from FCT – Fundação para a Ciência e Tecnologia, under the project: UIDB/04561/2020.

## Supporting Information

**S1 Appendix**.

**S2 Appendix. Model input parameters**.

**S1 Table. Proportion of B lineage influenza virus not included in the seasonal trivalent vaccines from 2010/11 to 2017/18 influenza seasons**.

**S2 Table. Description of the variables that take part of S1 Fig**.

**S3 Table. Input parameters**. Base case value; SD; parameters of the probability distribution.

**S4 Table. Probabilistic sensitivity analysis**. Assessing the 2015/16 influenza season-related uncertainty.

**S1 Fig. Probability, cost and QALY calculations by decision tree pathway for individuals aged** ≥**65 years old who received TIV**.

**S2 Fig. One-way sensitivity analysis of TIV effectiveness**. Variation between 34% and 74% with increments of 0.005.

**S3 Fig. One-way sensitivity analysis of QIV effectiveness**. Variation between 58% (TIV effectiveness) and 100% with increments of 0.005.

**S4 Fig. One-way sensitivity analysis of vaccination coverage rate**. Variation between 0% and 100% with increments of 0.05.

**S5 Fig. Probabilistic sensitivity analysis for TIV and QIV**. Scatter plot based on the 2015/16 influenza season.

