## Supplementary material for "Cost-effectiveness of quadrivalent versus trivalent inactivated influenza vaccines for the Portuguese elderly population": S1 Appendix

The cost-effectiveness analysis carried out in this paper relies on the computation of QALYs, which has become widely used in this type of studies. It is thus assumed that the major objective is to maximize health or health improvement across the population subject to resource constraints. The greatest advantage of QALY is the combination of changes in morbidity and mortality in a single indicator. The core concept of the conventional QALY is grounded in decision science and expected utility theory(61). In order to estimate the value attributed to the different health states by the Portuguese individuals (that is, to estimate the utilities), we used the health questionnaires developed by the EuroQol, with special emphasis to EQ-5D. Thus, the health utilities assigned to the different state of health under consideration are based on published studies that applied the EQ-5D instrument.

There are few studies published in Portugal about QALY weights calculation. Utilities associated with a healthy state were obtained from a study on EQ-5D Portuguese population norms(16). Data were stratified by age, but there was no stratum for people aged 65 or over. In the same way, no studies were found about disutilities associated with influenza. Such data were extracted from Hollmann et al.(17), where QALYs losses due to no hospitalized influenza and ILI without influenza confirmation were derived from individual outpatients QALYs losses. On the other hand, disutilities associated with hospitalizations due to influenza, pneumonia, respiratory disease and heart disease were derived from individual inpatient QALYs losses(17), based on the EQ-5D system.

QALYs were calculated annually, as the time horizon chosen was one year. In the pathways ending in the “death” state, the utility associated with a healthy state was divided by 2. This happens because no time of death is known, so it is assumed to have occurred in the middle of the year.
