## Supplementary material for "Cost-effectiveness of quadrivalent versus trivalent inactivated influenza vaccines for the Portuguese elderly population": S2 Appendix

**Model Input Parameters**

Most of the data were extracted from the Portuguese Hospital Morbidity Database (HMDB)(37), which is owned by the Central Administration of the Health System Portugal. It gathers information - on patient-level - about inpatient and outpatient administrative data as well as hospital episode data. The data extracted from the HMDB consist of admissions of patients aged 65 or above, with a principal diagnosis of influenza, pneumonia, respiratory disease, or heart disease in 2015. For hospitalizations due to pneumonia, respiratory disease, and heart disease, only subjects who experienced an episode of influenza as a secondary diagnosis were considered.

The total population aged ≥65-year-old was derived from an indicator released by Statistics Portugal, named *Resident population (Long series, start 1991 - No.) by Place of residence (NUTS - 2013), Sex and Age; Annual*(26). Data on vaccination coverage in 2015/2016 season were taken from the following report published by the (Portuguese) National Health Institute Dr. Ricardo Jorge (INSA): *Vacinação antigripal da população portuguesa na época 2015/2016*(62).

Trivalent inactivated vaccine effectiveness was established at 58% according to Demicheli et al(12). It was calculated as 1-RR, where RR (Risk Ratio) corresponds to the ratio between the proportions of patients who developed influenza in vaccinated and unvaccinated patients. The latter are patients who received placebo(12). Quadrivalent inactivated vaccine effectiveness was estimated through the study of the proportion of circulating B strains not included in TIV across several seasons, summarized in S1 Table. The methodology was adapted from Petri and Ruiz-Aragón(34). The data were retrieved from the Portuguese National Influenza Surveillance Programme (NISP) annual reports for seasons from 2010/11 to 2016/17(27–33). Data related to season 2017/18 were derived from the weekly report on influenza surveillance published by INSA(35). On average, 8.69% of the circulating strains correspond to mismatched TIV strains. The next step was to obtain the relative gain in effectiveness of using QIV instead of TIV, which was calculated as 8.69% × 0.38 = 3.30%, where 0.38 indicates the mean reduction in vaccine effectiveness due to B lineage mismatch(34).

***Computation of the Probabilities***

The probability of confirmed influenza for people aged ≥65 years was based on the NISP annual report for 2015/2016 season(31). Data on GP consultations were extracted from the National Health Service online platform named *Seasonal Health*(36). The number of patients (≥65y) who attended the GP consultation in the context of confirmed influenza, denoted by
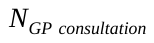
, is given by

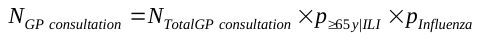

where
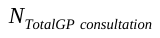
represents the total number of GP consultations for influenza, that is, the sum of GP consultations per week, from week 40 of 2015 to week 20 of 2016;
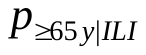
is the population proportion of patients aged ≥65 years who have ILI, obtained from the NISP annual report for 2015/2016 season(31);
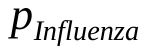
is the population proportion of patients with confirmed influenza obtained from the same report(31).

The probability of ILI, denoted by
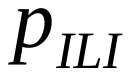
, takes thus the form

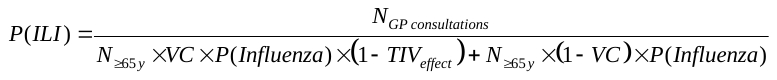

where
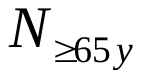
is the total number of individuals aged 65 and over in 2015, described in section “Input Parameters”; *VC*, the vaccination coverage rate, and
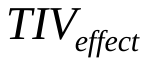
, TIV effectiveness, are also described in “Input Parameters”. It should be noted that the TIV effectiveness was considered in the calculation of ILI because in the 2015/16 influenza season only the TIV vaccine was administered.

Probabilities of hospitalization due to influenza, pneumonia, respiratory disease and heart disease as well as probabilities of death when hospitalized due to such conditions were obtained from the HMDB(37). Probability of death without influenza confirmed was based on the following indicators developed by Statistics Portugal: *Resident population (Long series, start 1991 - No.) by Place of residence (NUTS - 2013), Sex and Age; Annual* (26) and *Deaths (No.) by Place of residence (NUTS - 2013), Sex and Age; Annual*(38).

***Costs***

Cost of GP consultation and cost of vaccine administration were established by the Portuguese Government and can be found in the Ordinance No. 234/2015 of 7^th^ August, issued by the State Secretary for Health. All costs related to hospitalizations were calculated from the Portuguese Hospital Morbidity Database (HMDB)(37). The “episode cost” is obtained by the product of “relative weight”, “equivalent patient” and “base price”. The first term is available on the Price Table defined by the NHS for 2015(39); “equivalent patient” allows adjusting the “episode cost” according to the episode length, which begins with inpatient admission and ends with inpatient discharge; “inpatient episode” may be classified as normal, short or long stay according to the interquartile range of the respective Diagnosis-Related Group (DRG) (i.e., normal if it falls within the range, short if it is less than the lower bound and long if it is greater than the upper bound). Thus, each inpatient episode is converted into an equivalent patient considering the interquartile range defined for each DRG and the duration of inpatient episode. Finally, the inpatient base price was defined by the Central Administration of the Health System Portugal and it corresponds to 2,285€(39). Input costs of hospitalizations and deaths correspond to the average costs. It is important to note that the cost of death when hospitalized includes all costs related to the last hospitalization of the patient, so the cost of hospitalization is not considered in the pathways of the decision tree that leads to the “death” state.

Cost of Antiviral Treatment (AT) and cost of TIV for NHS were retrieved from online public contracts (available on the online platform: *BASE: Contratos Públicos Online*(40)). According to Orientation Nº 007/2015 from the Regulatory Decree Nº 14/2012 of 26^th^ January(41), article 2, paragraph 2(a), it was recommended the administration of the oseltamivir 75mg twice daily for 5 days. As a result, the total cost of 10 capsules was considered. Cost of AT was considered in all pathways with ILI. It is recommended to be administered at the earliest stage, preferably in the first 48 hours after the onset of the symptoms, and laboratory confirmation is not needed during influenza activity season. It was assumed that the cost of hospitalization and cost of death when hospitalized include the cost of antiviral treatment, so it was not added in such pathways.

Cost of TIV in 2015 was not available, so it was assumed to be the same as in 2016(42). As QIV was recently reimbursed by the NHS, the model was updated with QIV cost from 2019/2020 season, obtained from the online platform(43). It was applied a discount rate according to the average growth rate of the consumer price indexes from 2016 to 2019, which was retrieved from Statistics Portugal(44).
