## Supplementary material for "Cost-effectiveness of quadrivalent versus trivalent inactivated influenza vaccines for the Portuguese elderly population": S1 Table

**S1 Table. Proportion of B lineage influenza virus not included in the seasonal trivalent vaccines from 2010/11 to 2017/18 influenza seasons**.

| Season | Proportion of Influenza B | Proportion of B/Victoria | Proportion of B/Yamagata | B - lineage in TIV | Proportion of B lineage mismatch | SE | Source |
| --- | --- | --- | --- | --- | --- | --- | --- |
| 2010/11 | 43,20% | 42,70% | 0,50% | Victoria | 0,50% | 0,00022 | ^28^ |
| 2011/12 | 2,30% | 0,00% | 2,30% | Victoria | 2,30% | 0,00058 | ^29^ |
| 2012/13 | 51,30% | 1,80% | 49,50% | Yamagata | 1,80% | 0,00038 | ^30^ |
| 2013/14 | 0,80% | 0,10% | 0,70% | Yamagata | 0,10% | 0,00011 | ^31^ |
| 2014/15 | 0,36% | 0,00% | 0,36% | Yamagata | 0,00% | 0,00000 | ^32^ |
| 2015/16 | 8,30% | 7,80% | 0,50% | Yamagata | 7,80% | 0,00084 | ^33^ |
| 2016/17 | 0,20% | 0,20% | 0,00% | Victoria | 0,00% | 0,00000 | ^34^ |
| 2017/18 | 66,00% | 9,00% | 57,00% | Victoria | 57,00% | 0,00354 | ^35^ |
| 2010/11-2017/18 | 21,56% |  |  |  | 8,69% | 0,00213 |  |

SE: Standard Error
