## Supplementary material for "Cost-effectiveness of quadrivalent versus trivalent inactivated influenza vaccines for the Portuguese elderly population": S2 Table

**S2 Table. Description of the variables that take part of S1 Fig.**

| **Code** | **Description** | **Code** | **Description** |
| --- | --- | --- | --- |
| p_ili | Probability of Influenza-like-illness | c_death_hosp_RD | Cost of death when Hospitalization due to RD |
| p_conf_ili | Probability of Confirmed Influenza | c_hosp_HD | Cost of Hospitalization due to Heart Disease |
| p_gp | Probability of GP Consultation | c_death_hosp_HD | Cost of death when Hospitalization due to HD |
| p_hosp_inf | Probability of hospitalization due to Influenza | c_death | Cost of death |
| p_death_inf | Probability of death when Hosp. Influenza | c_ant_treat | Cost of antiviral treatment |
| p_hosp_pneu | Probability of hospitalization due to Pneumonia | c_tiv | Cost of trivalent vaccine |
| p_death_pneu | Probability of death due to Hosp. Pneumonia | c_qiv | Cost of quadrivalent vaccine |
| p_hosp_RD | Probability of hospitalization due to Respiratory Disease | c_vac_admin | Cost of vaccine administration |
| p_death_RD | Probability of death when Hosp. RD | u_ili_no_inf | Disutility associated with ILI without influenza confirmation |
| p_hosp_HD | Probability of hospitalization due to Heart Disease | u_inf_no_hosp | Disutility associated with no hospitalized influenza |
| p_death_HD | Probability of death when Hosp. HD | u_hosp_inf | Disutility associated with hospitalization due to influenza |
| p_death_no_inf | Probability of death when no confirmed influenza | u_hosp_pneu | Disutility associated with hospitalization due to pneumonia |
| c_ili_no_inf | Cost of ILI without influenza confirmation | u_hosp_RD | Disutility associated with hospitalization due to respiratory disease |
| c_GP | Cost of GP consultation | u_hosp_HD | Disutility associated with hospitalization due to heart disease |
| c_hosp_inf | Cost of Hospitalization due to Influenza | u_healthy | Utility associated with healthy population |
| c_death_inf | Cost of death when Hosp. Influenza | Pop | Population |
| c_hosp_pneu | Cost of Hospitalization due to Pneumonia | Coverage | Vaccination Coverage Rate |
| c_death_hosp_pneu | Cost of death when Hospitalization due to Pneumonia | tiv_effect | Trivalent Vaccine Effectiveness |
| c_hosp_RD | Cost of Hospitalization due to Respiratory Disease | qiv_effect | Quadrivalent Vaccine Effectiveness |
