## Supplementary material for "Cost-effectiveness of quadrivalent versus trivalent inactivated influenza vaccines for the Portuguese elderly population": S3 Table

**S3 Table. Input parameters.** Base case value; SD; parameters of the probability distribution.

| **Input Parameters** | **Base Case** | **SD** | **Distribution** | **Parameter 1** | **Parameter 2** |
| --- | --- | --- | --- | --- | --- |
| Probability of Confirmed Influenza | 0.278 | 0.037 | Beta | 39.72 | 103.28 |
| Probability of being ≥65years when ILI | 0.130 | 0.0095 | Beta | 162.87 | 1089.13 |
| Disutility associated with ILI without influenza confirmation | 0.009 | 0.0010 | Gamma | 77.79 | 0.00012 |
| Disutility associated with no hospitalized influenza | 0.009 | 0.0010 | Gamma | 77.79 | 0.00012 |
| Disutility associated with hospitalization due to influenza | 0.031 | 0.0031 | Gamma | 102.55 | 0.00030 |
| Disutility associated with hospitalization due to pneumonia | 0.031 | 0.0031 | Gamma | 102.55 | 0.00030 |
| Disutility associated with hospitalization due to respiratory disease | 0.031 | 0.0031 | Gamma | 102.55 | 0.00030 |
| Disutility associated with hospitalization due to heart disease | 0.031 | 0.0031 | Gamma | 102.55 | 0.00030 |
| Utility associated with healthy population | 0.625 | 0.0163 | Gamma | 525.41 | 0.00071 |
| Vaccination coverage rate | 0.501 | 0.041 | Beta | 74.68 | 74.38 |
| Proportion of B lineage viruses not included in TIV | 0.087 | 0.002 | Beta | 15.77 | 165.81 |
| Relative Risk  (1-Trivalent Vaccine Effectiveness) | 0.421 | 0.228 | Lognormal | -0.865 | 0.228 |

SD: Standard Deviation
