## Supplementary material for "Cost-effectiveness of quadrivalent versus trivalent inactivated influenza vaccines for the Portuguese elderly population": S4 Table

**S4 Table. Probabilistic sensitivity analysis.** Assessing the 2015/16 influenza season-related uncertainty.

|  | **TIV** | **QIV** | **Difference (QIV-TIV)** |
| --- | --- | --- | --- |
| Events |  |  |  |
| GP Consultations | 2,636.73 | 2,602.02 | -34.71 |
| Hospitalizations due to Influenza | 239.00 | 235.85 | -3.15 |
| Deaths due to Influenza Hospitalization | 19.00 | 18.75 | -0.25 |
| Hospitalizations due to Pneumonia | 21.00 | 20.72 | -0.28 |
| Deaths due to Pneumonia Hospitalization | 8.00 | 7.89 | -0.11 |
| Hospitalizations due to RD | 40.00 | 39.47 | -0.53 |
| Deaths due to RD Hospitalization | 2.00 | 1.97 | -0.03 |
| Hospitalizations due to HD | 73.00 | 72.04 | -0.96 |
| Deaths due to HD Hospitalization | 8.00 | 7.89 | -0.11 |
| Vaccine Doses | 1,037,847 | 1,037,847 | 0.00 |
| Costs |  |  |  |
| GP Consultations | 81,739 € | 80,663 € | -1,076 € |
| Hospitalizations due to Influenza | 710,476 € | 701,447 € | -9,029 € |
| Deaths due to Influenza Hospitalization | 121,553 € | 119,977 € | -1,576 € |
| Hospitalizations due to Pneumonia | 128,214 € | 126,553 € | -1,661 € |
| Deaths due to Pneumonia Hospitalization | 90,401 € | 89,220 € | -1,181 € |
| Hospitalizations due to RD | 99,127 € | 97,877 € | -1,250 € |
| Deaths due to RD Hospitalization | 2,731 € | 2,698 € | -33 € |
| Hospitalizations due to HD | 256,052 € | 252,779 € | -3,273 € |
| Deaths due to HD Hospitalization | 62,303 € | 61,493 € | -810 € |
| Vaccine Doses | 2,673,494 € | 7,983,576 € | 5,310,082 € |
| Total | 8,029,692 € | 13,323,148 € | 5,293,456 € |
| CI lower bound | 6,904,148 € | 11,292,400 € | 4,375,217 € |
| CI upper bound | 9,018,064 € | 15,125,063 € | 6,088,637 € |
| QALYs |  |  |  |
| Total | 1,265,457.89 | 1,265,458.09 | 0.20 |
| CI lower bound | 1,197,368.63 | 1,197,368.75 | 0.06 |
| CI upper bound | 1,328,396.10 | 1,328,396.36 | 0.43 |
| ICER (€/QALY) |  |  | 34,501,793 |
| CI lower bound |  |  | 13,052,665 |
| CI upper bound |  |  | 87,418,632 |
